# Single-cell multiomics identifies both shared and unique features of immune dysfunction in Parkinson’s disease and inflammatory bowel disease colon, plasma and stool

**DOI:** 10.1101/2025.04.29.651228

**Authors:** MacKenzie L. Bolen, Marc Buendia, Ji Shi, Hannah Staley, Jennifer M. Kachergus, Philip A. Efron, Gwoncheol Park, Ravinder Nagpal, Stephan D. Alvarez, Qing-Shan Xue, Nikolaus R. McFarland, Ellen M. Zimmermann, Christopher E. Forsmark, Kelly B. Menees, Azucena Salas, E. Aubrey Thompson, Malú Gámez Tansey

## Abstract

Parkinson’s disease (PD) is the fastest growing neurodegenerative disease in the world^1^. Gastrointestinal (GI) dysfunction can occur decades before motor impairments and in up to 80% of individuals living with PD^2,3,4^. We investigated peripheral relationships that may underlie mechanisms along the gut-blood axis that contribute to PD pathogenesis. Single-cell multiomic spatial molecular imaging (SMI) of colonic tissue localized inflammatory injury within epithelial cells that appear to be associated with iron mishandling in both inflammatory bowel disease (IBD) and PD biosamples. We found that both the single-cell SMI of RNA and protein revealed parallel cross-modal dysregulation in the gut epithelium, in both IBD and PD biosamples. These data are accompanied by plasma (PD) and stool (IBD) protein depletion of CCL22. Our findings suggest iron mishandling along the gut barrier likely contributes to systemic inflammation, which may be the catalyst that primes circulating immune cells to body-first PD pathogenesis.

## Introduction

The rapid growth in Parkinson’s disease (PD) diagnoses in the last decade is likely due to an increase in environmental risk factors such as unhealthy diets, pesticides, and pollutants related to industrialization that have all been implicated in PD risk^5^. However, PD is a spectrum disorder, meaning the stereotypical bradykinesia or tremor used to diagnose this neurodegenerative disease often occurs up to 20 years after nonmotor symptoms including anosmia, sleep disturbances, and gastrointestinal (GI) dysfunction^2,6^. The delayed timeline from nonmotor symptoms to motor symptom presentation has prompted intense investigations into the possibility that environmental barrier sites outside of the brain could be the nexus of PD pathogenesis. One such peripheral site is the GI tract and its role in body-first PD^7^. Gut dysfunction, such as constipation, often occurs decades prior to motor deficits in up to 80% of those living with PD^8^. It is also known that those living with inflammatory bowel disease (IBD), the most common subtype of gastrointestinal (GI) dysfunction, have a 30% increased risk of developing PD^9^. Importantly, not all individuals with IBD develop PD, and vice versa, but individuals with IBD may be predisposed to development of body-first PD as opposed to brain-first PD^10^.

The molecular basis connecting gut dysfunction in IBD to increased risk for PD is still not understood. A primary issue with investigating the gut-brain axis is the heterogeneity in biological features and validated clinical symptom presentation of early phase gut dysfunction, such as in IBD^11,12^. Therefore, to our knowledge, we have evaluated the largest sigmoid colonic biopsy cohort to date (IBD, in remission N = 13; PD N = 12; NHC N = 9) to appreciate the spatial single-cell targeted transcriptome and proteomic landscape; and thus, represent as much of this heterogeneous population as possible.

It is known that the epithelial lining of the gut separates the host’s intestinal lamina propria where gut-resident immune cells reside from bacterial-derived products and pathogens; when this barrier is breeched or compromised, there is increased mucosal permeability to harmful antigens^13^. IBD (including both ulcerative colitis and Crohn’s disease) is associated with chronic relapsing and often transmural inflammation of the intestine^14^. This inflammatory cascade drives the breakdown of the intestinal lining and contributes to development of a “leaky gut” and uncontrolled release of chemokines and cytokines that prompt release of apoptotic and necrotic cell factors^15^. A leaky gut drives a proinflammatory immune response by increasing mucosal immune cell engagement with antigens present in the gut and allows for microbial products present in the gut to leak into deeper layers of the intestine within the peritoneal cavity^16^.

This relapsing-remitting cycle of cytokines drives the activation of the innate and adaptive immune systems^17^, leading to a feedforward mechanism of further cytokine and proinflammatory product release. Of note, those living with IBD who are prescribed anti-TNF biologics, common immune-targeted brain impermeant anti-inflammatory treatments, have a ∼75% decreased risk of developing PD compared to those living with IBD not on anti-TNF medication^18^. These epidemiological associations and anti-inflammatory intervention outcomes suggest that the peripheral immune system is an important messenger in the gut-brain axis in the context of PD.

There has yet to be an identified biomarker(s) indicating a molecular mechanism that links these two diseases of inflammation, likely due to the breadth of existing literature that places a heavy emphasis on a brain-centric approach to neurodegenerative disease. We, as well as others^19,20^, hypothesize that PD pathogenesis may be initiated in the gut decades prior to the development of classic PD motor phenotypes in part due to disruptions in central-peripheral neuroimmune crosstalk^21^.

We interrogated the unique versus shared cellular and molecular features of mechanisms associated with dysregulated gut immunity and gut inflammation in IBD, PD, and neurologically healthy controls (NHCs). We employed NanoString CosMx^TM^ Spatial Molecular imaging (SMI) to conduct multiplexed spatial imaging of both the targeted transcriptome and proteome to profile the heterogeneous mosaic of the colon and immune-interacting cellular partners within the colonic microenvironment. The use of multiplexed protein immunoassays of biospecimens at additional peripheral sites (plasma and stool) allowed for additional in-depth and cross-correlational analyses of the gut-blood axis in individuals with IBD (in remission), PD, or NHCs. Our findings provide a foundational open-access data repository that can be leveraged to identify peripheral biomarkers of PD risk in the gut-blood axis (accessible via Zenodo: https://doi.org/10.5281/zenodo.14851478 (RNA) https://doi.org/10.5281/zenodo.14851272 (protein).

## Results

### Clustering analysis identifies cell-type abundance and mapping of spatial organization unique to PD and IBD in sigmoid colonic biopsies

Colonic biopsies from PD (*N* =12), IBD in remission (*N* = 13, including *n* = 9 diagnosed Crohn’s disease, *n* = 3 diagnosed ulcerative colitis and *n* = 1 IBD with no defined diagnosis) and NHCs (*N* = 9) were collected to interrogate intrinsic immune and inflammatory features within each compartment (Supplemental Table 1; Supplemental Fig. 1). It is important to note that all biopsies collected were deemed endoscopically noninflamed by a gastroenterologist at the time of collection, due to this classification all individuals diagnosed with ulcerative colitis or Crohn’s disease were pooled together into one IBD cohort. Compartments consisting of primary cell types (epithelial, B and plasma cells, stromal, T cells, myeloid cells) (Fig. 1D; Supplemental Table 2; Supplemental Fig. 2) were isolated *in silico* to generate higher resolution of refined subpopulations (Fig. 1B; Supplemental Fig. 3 and Supplemental Fig. 4). Annotated refined populations were then digitally mapped onto tissue. The spatial distribution of relevant cell types appeared to have the expected characteristics of sigmoid colon (Fig. 1C) and was unique to IBD, PD or NHC (Fig. 1D). Additionally, refined cell type abundance differed by disease (Fig. 1E). However, there was a significant increase in the abundance of colonocytes in both IBD and PD patients (Fig. 1E); but interestingly, there was a significant decrease in CD4+ T cells in those living with IBD and an increase in CD8+ T cells in those living with PD, compared to NHC (Fig. 1E). A subsequent slide was then analyzed for protein regulation, however only to the depth of subpopulations. Colonic biopsies from either those living with IBD (in remission) or PD displayed a significant enrichment in both proteins identified subpopulations of epithelial and T cells (Fig. 6C).

**Figure 1.**
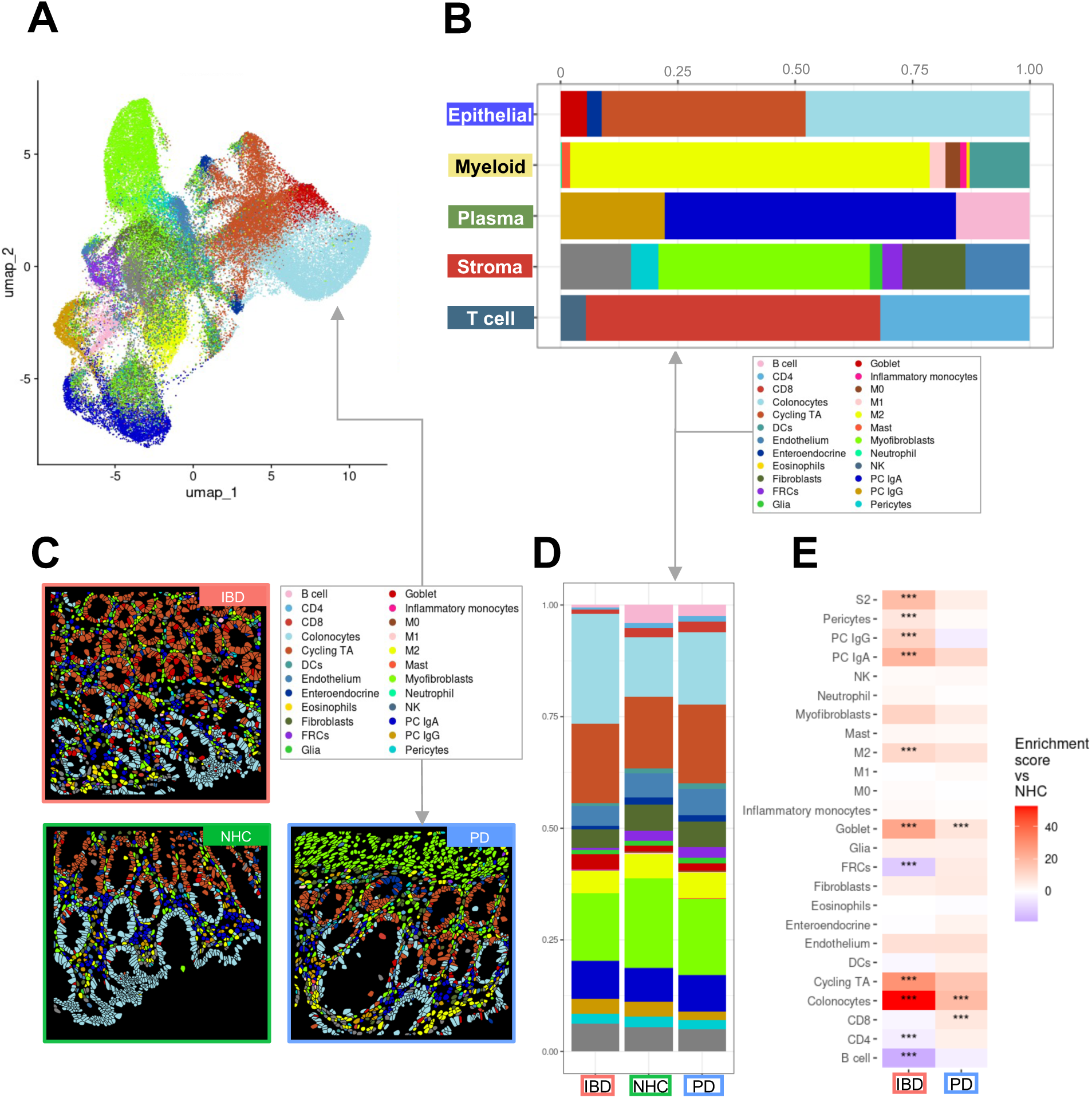
Spatial molecular imaging (SMI) enables deep immune-phenotyping of sigmoid colon biopsies from those living with inflammatory bowel disease (IBD) and Parkinson’s disease (PD). A *In situ* uniform manifold approximation and projection (UMAP) clustering of all cells identified within colonic tissue samples (N = 13 IBD, N = 12 PD, N = 9 NHC). Cells are colored by all annotated refined clusters identified (epithelial, myeloid, plasma, stroma and T cells). Cells were only included if the probability of being correctly annotated within the respective cluster was greater than 0.75. **B** Bar plots of SMI data are colored by group cluster identifier and shaded by refined subset population within each primary cluster. Each bar plot represents the proportion of refined subpopulation in each primary cluster of cells. **C** CosMx^TM^ SMI visualization indicating cell-type specific spatial distribution of all refined populations annotated. **D** Bar plot representation of refined subcluster proportion of cells by disease group. Source data my be found in the Source Data file provided in supplement. **E** Heatmap depicting cell-type abundance by IBD (red, x-axis) or PD (blue, x-axis) as compared to neurologically healthy control (NHC). Statistical analysis completed by Chi-Square (χ²) test with p value adjustment using Benjamini–Yekutieli test. Only results with a fold change greater than 1.5 or less than 0.66 are considered significant after adjustment. * = adjusted p-value less than 0.05, ** = adjusted p-value less than 0.01, ***= adjusted p-value less than 0.001.

### Differential gene expression identifies inflammatory pathways associated with immune signaling dysfunction in the colon in both PD and IBD

Dysregulation of iron absorption is a known outcome of PD in the brain^22^ and can be a marker of inflammation and autoimmune dysfunction in circulating peripheral immune cells^23^. The role gut inflammation plays in peripheral iron handling and how this inflammatory catalyst may be linked to PD has yet to be unraveled. Volcano plots depicting differentially expressed genes (DEGs) comparing epithelial cell clusters in PD vs NHC revealed enriched markers of inflammation in both IBD and PD colonic biopsies (Fig. 2A & Fig. 2B). Here, we observed a significant enrichment of one of the genes that encode for the major iron storage protein ferritin, the ferritin heavy chain 1 (*FTH1*), in pooled epithelial IBD biopsies (Fig. 2A); and in pooled epithelial PD biopsies depletion of *FTH1* and the gene that encodes for ferroportin (*SLC40A1),* the protein that drives ferritin export (Fig. 2B), relative to NHC biopsies. Visual spatial distribution of cells annotated as epithelial (orange) and *FTH1* single-cell gene expression (yellow) revealed significantly enriched *FTH1* expression in epithelial cells in those living with IBD in remission (Fig. 2Ai, Fig. 2Aii, Fig. 2Bi & Fig. 2Bii). In colonocytes from IBD biopsies, we observed a depletion in protein exporter gene *SLC40A1* and enrichment in genes associated with cell proliferation and migration including *MTOR* (mammalian target of rapamycin), *LGALS3* (lectin, galactoside-binding, soluble, 3) and *TIMP1* (TIMP metallopeptidase inhibitor 1) (Fig. 2C). In colonocytes from PD biopsies, *FTH1* and *SLC40A1* were significantly depleted (Fig. 2D). *MT2A*, a critical marker of metal homeostasis and immune regulation, was depleted in both IBD and PD colonocytes as compared to NHC (Fig. 2C & 2D). Relative density plot of total *FTH1* expression by disease and all refined cell types indicated that colonocytes contained the highest abundance of *FTH1* in the sigmoid colon regardless of disease status (Fig. 2E). Similarly, *SLC40A1* relative density plot revealed high abundance in colonocytes, and M2 macrophages and neutrophils in a disease-specific manner (Fig. 2F). Gene ontology (GO) biological process (BP), cellular component (CC) and molecular function (MF) analysis revealed several cellular mechanisms and resulting products that were significantly down- or up-regulated (Fig. 3). IBD epithelial cells displayed a marked decrease in expression of genes involved in cell chemotaxis, cytokine activity, leukocyte migration (Fig. 3A: blue); and a significant increase in genes involved in ameboid cell migration, response to extracellular stimulus and oxygen levels, as well as inflammatory responses (Fig. 3A:red). Epithelial cells from those living with PD displayed enrichment in GO pathways associated with neurogenesis, glial cell development and gliogenesis, as compared to NHC (Fig. 3B: red). IBD colonocyte GO pathways indicated depletion in chemotaxis signaling, leukocyte proliferation, and cytokine activation with an increase in regulation of response to external stimulus and ameboid cell migration (Fig. 3C).

**Figure 2.**
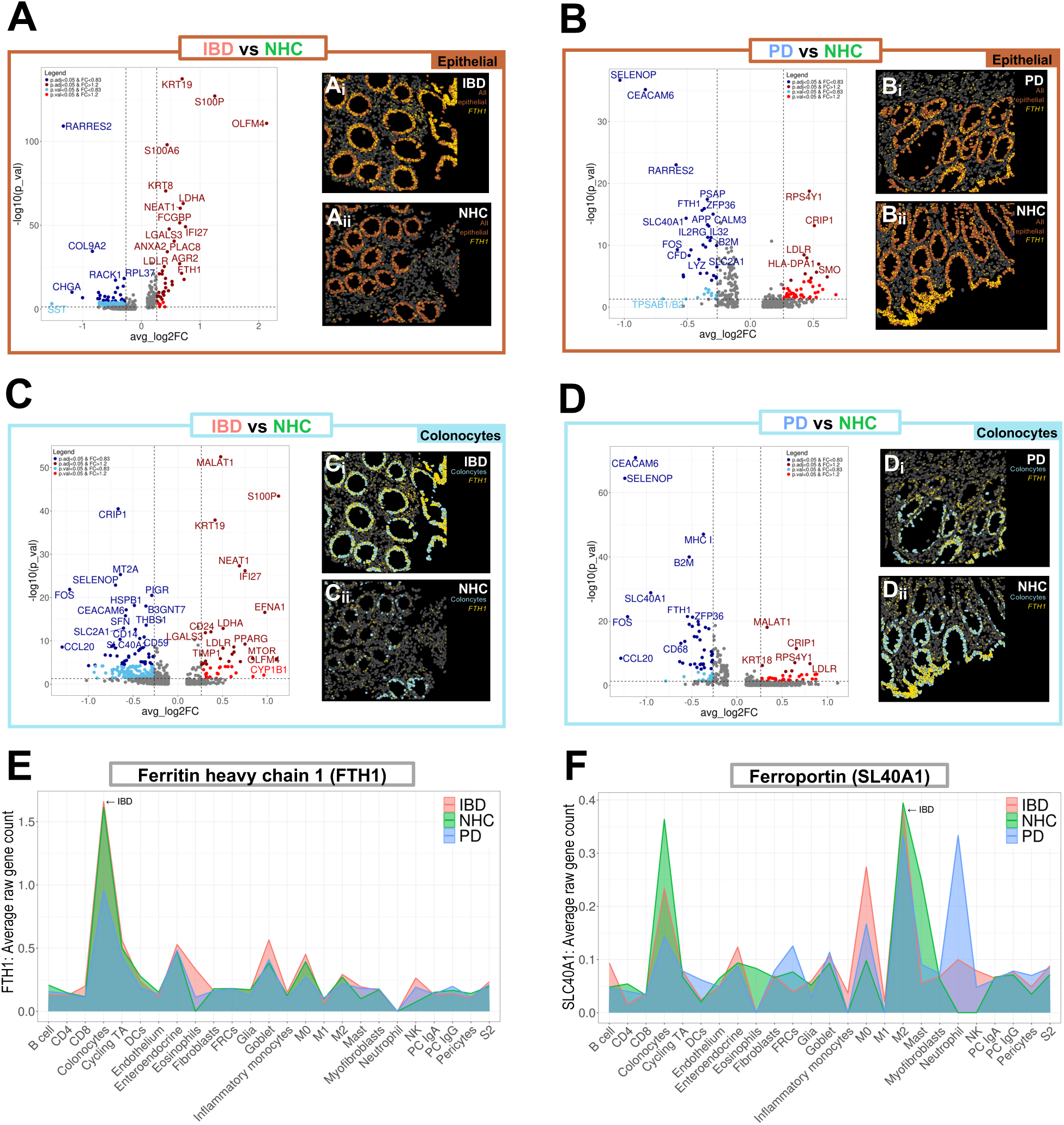
Epithelial gene profiles reveal enhanced inflammatory regulation in IBD and reduced inflammatory regulation in PD colonic biopsies. A,B,C,D Volcano plots of differential gene expression between IBD (red), PD (blue) or NHC (green) in all epithelial cells (brown box) or colonocytes (light blue box). Light blue indicates p_val < 0.05 & FC < 0.83, dark blue indicates p_adj_val < 0.05 & FC < 0.83, light red indicates p-value < 0.05 and dark red indicates p adjusted-value < 0.05 & FC > 1.2. A,B,C,D (i and ii) Visual spatial representation of one representative FOV from each disease group of epithelial cells (brown) or colonocytes (light blue) with gene counts of ferritin heavy chain 1 (*FTH1*) spatially resolved in yellow. E,F Cell-type specific density plot by average raw gene count of ferritin heavy chain 1 (*FTH1*) or ferroportin (*SLC40A1*) by disease group (IBD = red, NHC = green, PD = blue).

**Figure 3.**
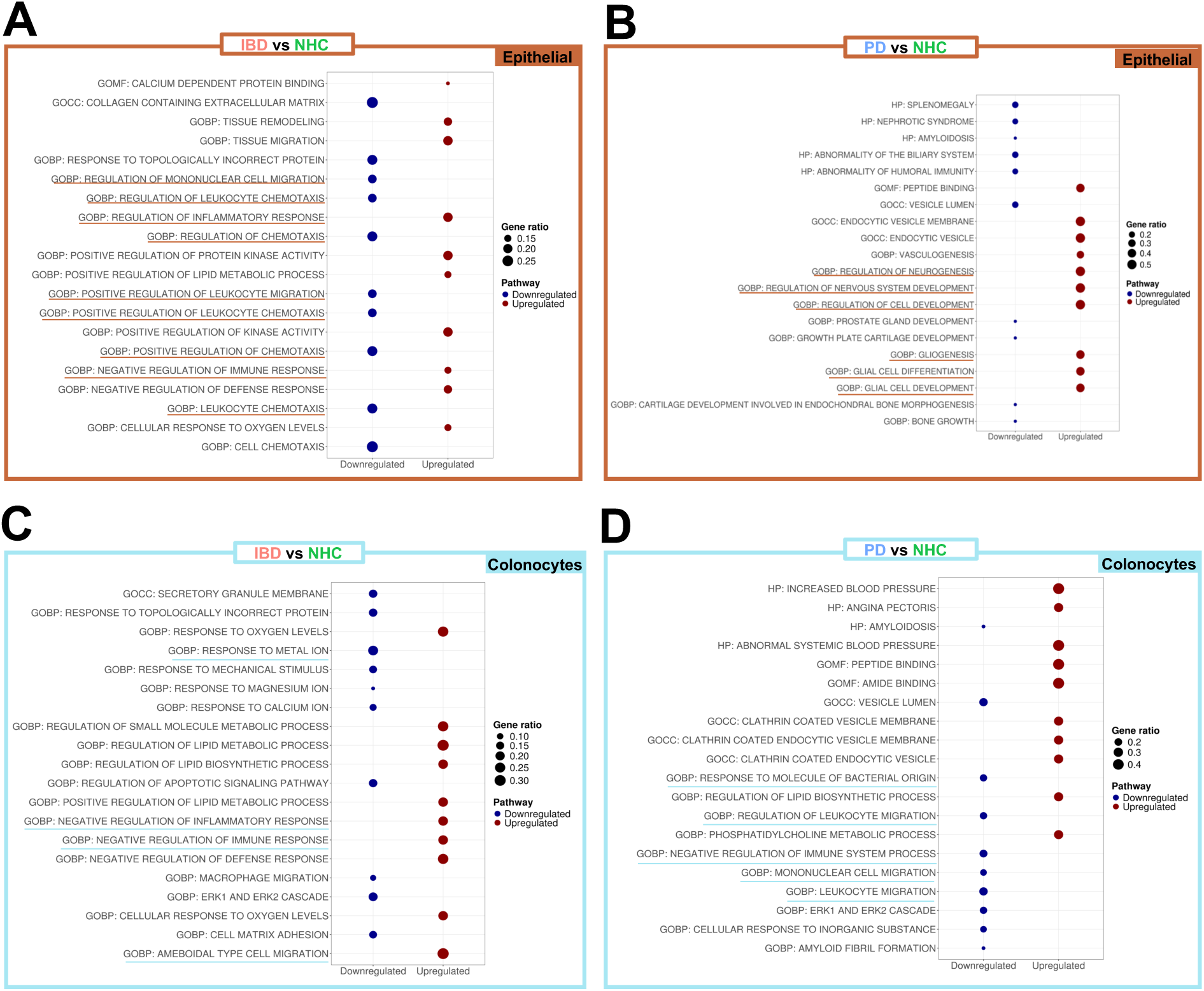
Colonocytes gene ontology pathway profile indicate enhanced inflammatory regulation in IBD and reduced inflammatory regulation in PD relative to NHC. **A,B,C,D.** Gene Ontology (GO) Human Category 5 specific pathways are compared between IBD, PD and NHC within epithelial (brown box) or colonocyte blue bow) populations to understand the mechanism behind broader pathway implications. Top 10 upregulated and downregulated pathways are displayed, ranked via most significant adjusted significance value (adjusted via false discovery rate, Benjamini–Yekutieli procedure). BP = biological process, MF = molecular function and CC = cellular component.

Additionally, colonocytes from those living with IBD but are in remission revealed a significant increase in gene expression involved in the negative regulation inflammatory and general immune response (Fig. 3C: red). GO pathway analysis of PD colonocytes revealed significant depletion in genes involved with cell adhesion, T cell migration, and myeloid differentiation (Fig. 3D: blue). Additional GO pathways evaluating thematic gene enrichment of PD colonocytes revealed significant enrichment in genes associated with response to external stimulus, T cell activation and leukocyte proliferation, with a decrease in migratory regulation and chemokine activity (Fig. 3D). Collectively, these data suggested a dampened immune activation status characterized by dampened cytokine expression in colonic leukocytes (i.e., neutrophils, lymphocytes, monocytes, eosinophils and basophils) (Fig. 3). These data suggested the presence of an inflammatory phenotype characterized by deficient expression of genes that promote and sustain immune-cell recruitment at the level of the sigmoid colon in those living with PD. Several other pathways appeared within GO pathway analysis as significantly up-regulated in IBD and down-regulated in PD (Fig. 3), however they are not discussed herein in great detail due to being outside of the scope of this manuscript.

### Spatial transcriptomics analysis reveals dysregulation in cell-cell crosstalk and neighborhoods at the level of the sigmoid colon in both IBD and PD

When assessing cell-cell communication, SCOTIA was used to generate chord plots and leverage ligand-receptor spatial analysis between source and target cells. Colonocytes positive for *FTH1* (green) or colonocytes negative for *FTH1* (blue) from individuals living with IBD in remission, PD, or NHC were assessed based on frequency of ligand-receptor interactions with other cells (Fig. 4A). Number of colonocyte cell-cell interactions were then quantified, wherein we found that there were significantly more interactions between *FTH1-*positive (*FTH1+)* cells and other cell types in IBD and PD colonic biopsies, but not NHC (Fig. 4B). Spatial connection maps displayed representative FOVs of colonocyte communication to *FTH1*+ cells (red) and colonocyte communication to *FTH1*-negative (*FTH1-)* cells (blue) (Fig. 4Ci-Cii). Those living with IBD and PD had significantly more *FTH1+* colonocyte ligand-receptor interactions with other cells than *FTH1*+ colonocytes from NHC (Fig. 4A; Fig. 4C-Ci). Additionally, colonocytes that are *FTH1*+ in those living with IBD that are in remission displayed significant depletion in *MTOR* and *LGALS3* (Fig. 4D). A traditional homeostatic marker of the mucosa, *MALAT1* (mucosal homeostatic regulator metastasis associated in lung adenocarcinoma transcript 1) ^24^ was depleted and *CRIP1* (cysteine-rich protein 1)^25^, which likely plays a role in immune cell activation, was significantly enriched; both suggesting aberrant mucosal homeostasis. Colonocytes that are *FTH1+* in those living with PD displayed an enrichment in *MHCI,* involved in antigen presentation to cytotoxic T cells^26^(Fig. 4Di). Both colonocytes positive for *FTH1* in those living with IBD in remission or those living with PD displayed a significant enrichment of *CEACAM6* (carcinoembryonic antigen–related cell adhesion molecule 6) which acts as a primary receptor for invasive *Escherichia coli* (*E. Coli*) colonization^27^ and *CCL22* (C-C motif ligand 22), a common chemotactic indicator of intestinal inflammation also known as *MDC* (macrophage-derived chemokine)^28^.

**Figure 4.**
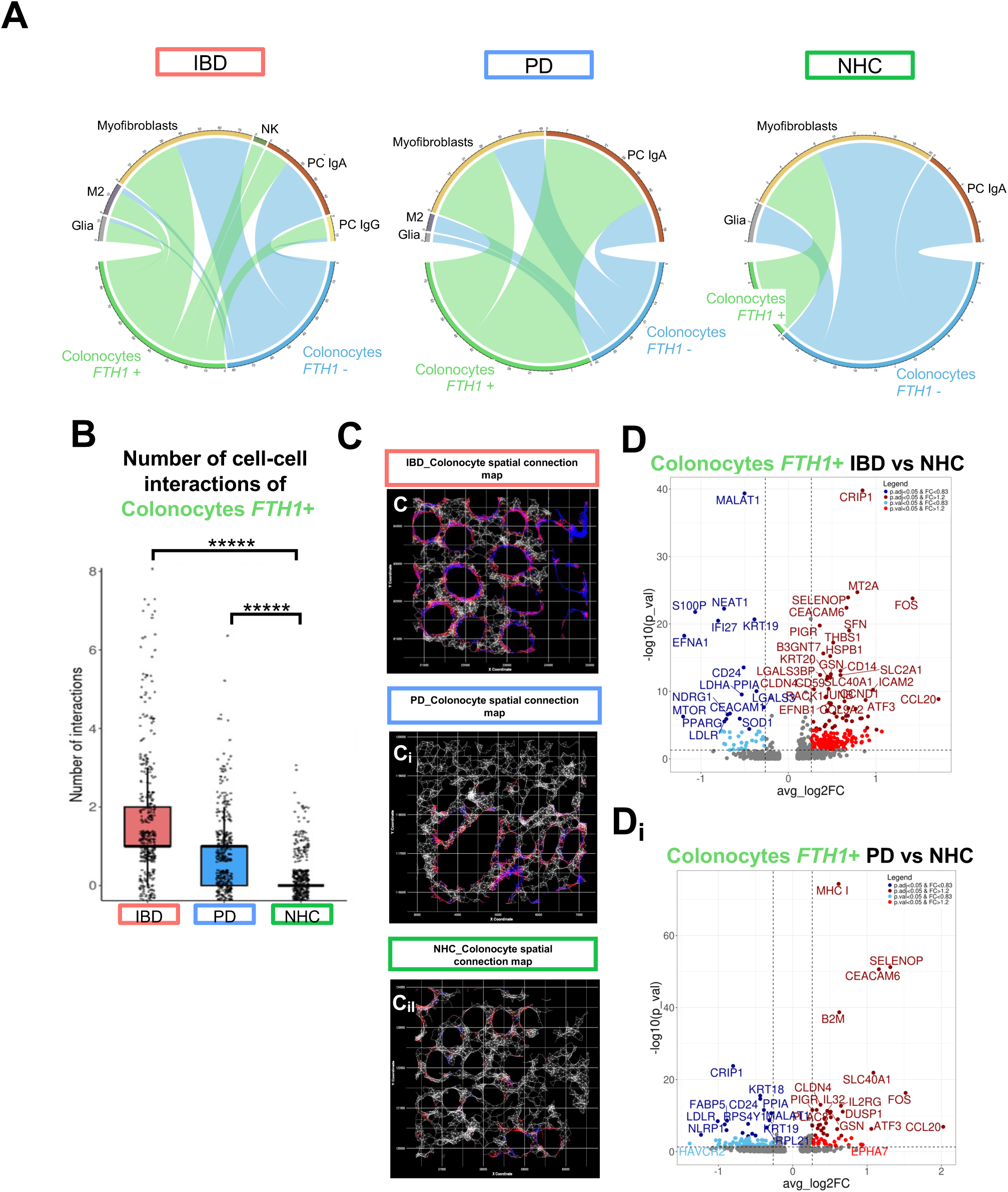
Spatial distribution of immune cells differs between IBD and PD relative to NHC. A-Aii Cell-cell communication of colonocytes either positive for ferritin heavy chain 1 (*FTH1*) (green) or negative for *FTH1* (blue) ligand-receptor interactions to all other refined cell populations based on disease group, generated via SCOTIA ligand receptor analysis. **B** Bar plots depicting number of significant interactions between the number of colonocytes that are *FTH1+* interacting with other cells by disease group. **C-Cii** Gene spatial connection map, white = all communication between all cells within 25uM radius, red = colonocytes connection to a ferritin heavy chain 1 (*FTH1*) positive cell, blue = colonocyte communication to *FTH1* negative cell. **D-Di** Volcano plots of differential gene expression between IBD colonocytes positive for *FTH1*, as compared to NHC and PD colonocytes positive for *FTH1*, as compared to NHC. Light blue indicates p_val < 0.05 & FC < 0.83, dark blue indicates p_adj_val < 0.05 & FC < 0.83, light red indicates p-value < 0.05 and dark red indicates p adjusted-value < 0.05 & FC > 1.2.

### Inflammatory factor immunoassays reveal a dysfunctional immune phenotype in both IBD and PD cohorts relative to NHC

One critical function of cytokines and chemokines is to recruit in adaptive immune cells to sites of injury^29^. Multiplexed immunoassays on the MesoScale Discovery (MSD) platform were leveraged to assess cytokine and chemokine signaling in the blood and stool from the same cohort of participants. It is important to note that the stool and blood biosample cohorts are slightly larger and not identical to the colonic biopsies due to sample availability. C-C motif chemokine 22 (CCL22), also known as macrophage-derived chemokine (MDC), displayed a significant DC-released T-cell recruitment protein, and was significantly depleted in plasma from those living with PD as compared to NHC (Fig. 5A). A linear regression analysis identified a significant negative relationship between CCL22 protein abundance and PD disease duration where individuals with longer disease duration had reduced levels of CCL22 (R^2^ = 0.32, *p*= 0.03) (Fig. 5Ai). Linear regressions assessing the relationship between CCL22 content and ferritin abundance separated by disease group revealed a substantial relationship in IBD (R^2^ = 0.60, *p*= 0.0020) and PD (R^2^ = 0.51, *p*= 0.006) plasma samples, where individuals with low CCL22 also had low ferritin but this relationship did not exist in NHC (R^2^ = 0.19, *p*= 0.21) (Fig. 5Aii). CCL22 total protein abundance was also measured in stool samples from the same participants (Fig. 5B). CCL22 was significantly depleted in stool samples from those living with IBD as compared to NHC (Fig. 5B). Linear regression analysis did not identify any relationship between CCL22 protein abundance and PD disease duration (R^2^ = 0.21, *p*= 0.09) (Fig. 5Bi). Linear regressions assessing a relationship between CCL22 content and ferritin abundance parsed apart by disease group revealed a significant relationship in IBD (R^2^ = 0.56, *p*= 0.0020) and PD (R^2^ = 0.38, *p*= 0.014) and NHC (R^2^ = 0.42, *p*= 0.042) where individuals with low CCL22 also had low ferritin across all participant groups (Fig. 5Bii). The full panel of cytokine and chemokines on the MSD is listed within the Methods.

**Figure 5.**
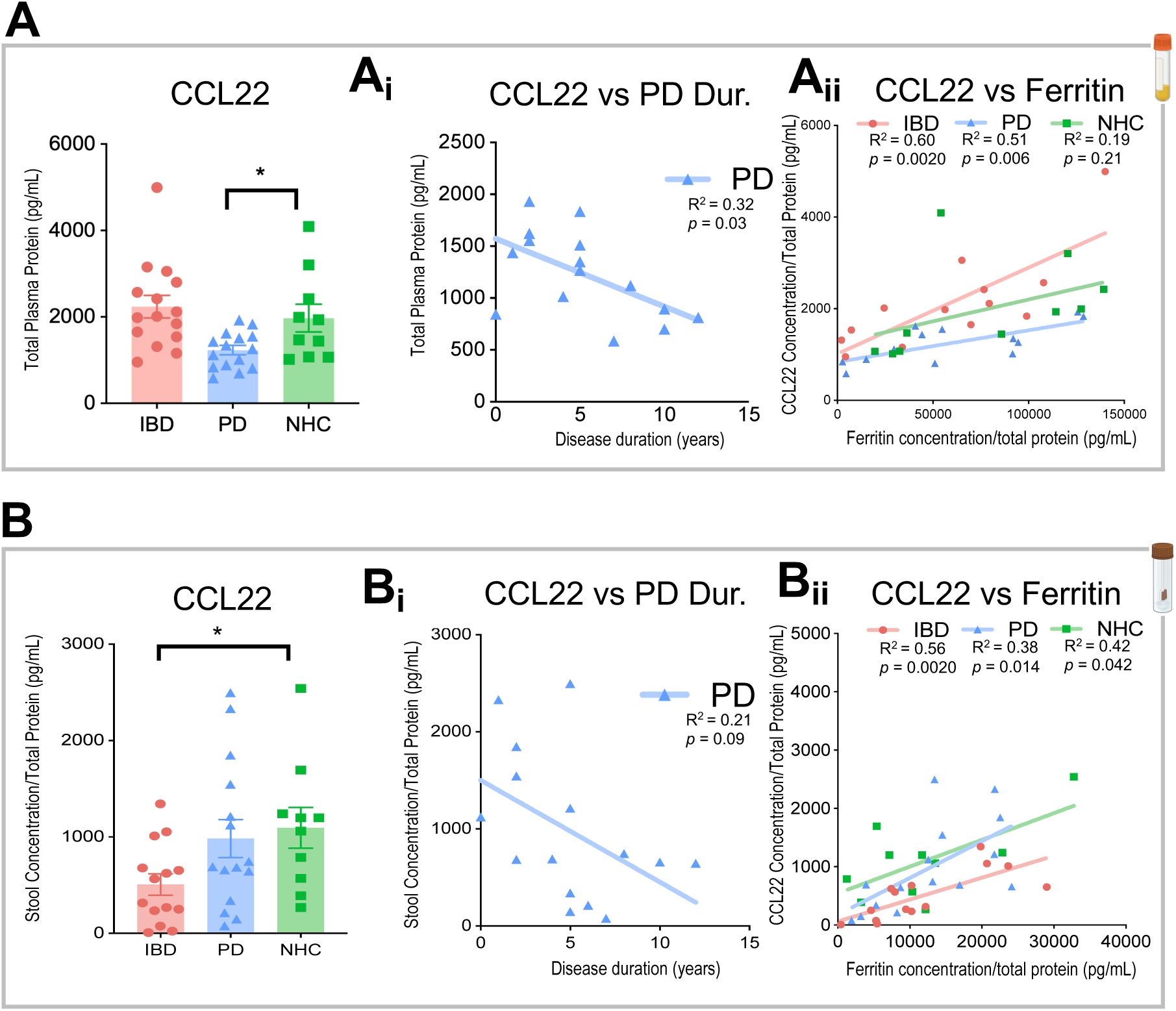
Metabolic analyses reveal alterations in markers of chemotaxis in plasma and stool in individuals living with IBD or PD, as compared to NHC. A-Aii In plasma samples (N = 17 IBD, N = 17 PD, N = 14 NHC), C-C motif chemokine ligand 22 (CCL22) is significantly depleted in PD as compared to NHC. There is no significant change observed between IBD and NHC. **B-Bii** In stool samples (N = 15 IBD, N = 15 PD, N = 10 NHC),, CCL22 is significantly depleted in IBD as compared to NHC. There is no significant change observed between PD and NHC. Linear regressions identify a significant correlation between increase in PD disease duration (DD) and decrease in CCL22 in plasma from those living with PD. Linear regression also displays a significant relationship with increase in CCL22 load and increase in ferritin load in both IBD and PD patients that does not appear in NHC (* = P<0.05). * = p-value less than 0.05, ** = p-value less than 0.01, ***= p-value less than 0.001, **** = p-value less than 0.0001.

### Single-cell targeted proteomic analysis successfully integrates with scRNA-seq and depicts multimodal immune dysregulation across multiple biological outputs of human colonic biopsies

To obtain the highest quality cell-type annotation of the targeted CosMx*^TM^* SMI proteomics data set, we integrated an existing single-cell RNA sequencing (scRNA-seq) dataset of human intestine biopsies^12^ using MaxFuse^30^ (FOV visualization Fig. 6A; Fig.6B; Supplemental Fig. 5). Weak links between protein and RNA overlay were chosen to generate UMAP clustering of 5 cell subpopulations (T cells, plasma cells, fibroblasts, epithelial cells and B cells) (Fig. 6Bi). Additionally, abundance analysis identified significant enrichment of epithelial, plasma and T cells in those living with PD or IBD, as compared to NHC (Fig. 6C). To interrogate the cell-cell communication, SCOTIA ligand-receptor analysis was leveraged to identify potential cell-type specific interactions. This analysis revealed an enrichment in communication between T cells in both those living with IBD or PD to epithelial cells as compared to NHC (Fig. 6D). We then analyzed immune checkpoint markers known to regulate immune responses in the context of inflammation and cancer^31^ and found a significant difference in PD-1 as well as CTLA4, GITR, GZMA, Ki-67, LAG3 and TIM3 in T cells housed in the colon of individuals living with PD, as compared to NHC (Fig. 6E). In contrast, T cells from those living with IBD in remission only displayed a significant difference in PD-1, GZMA, LAG3 and TIM3 (Fig.6E).

**Figure 6.**
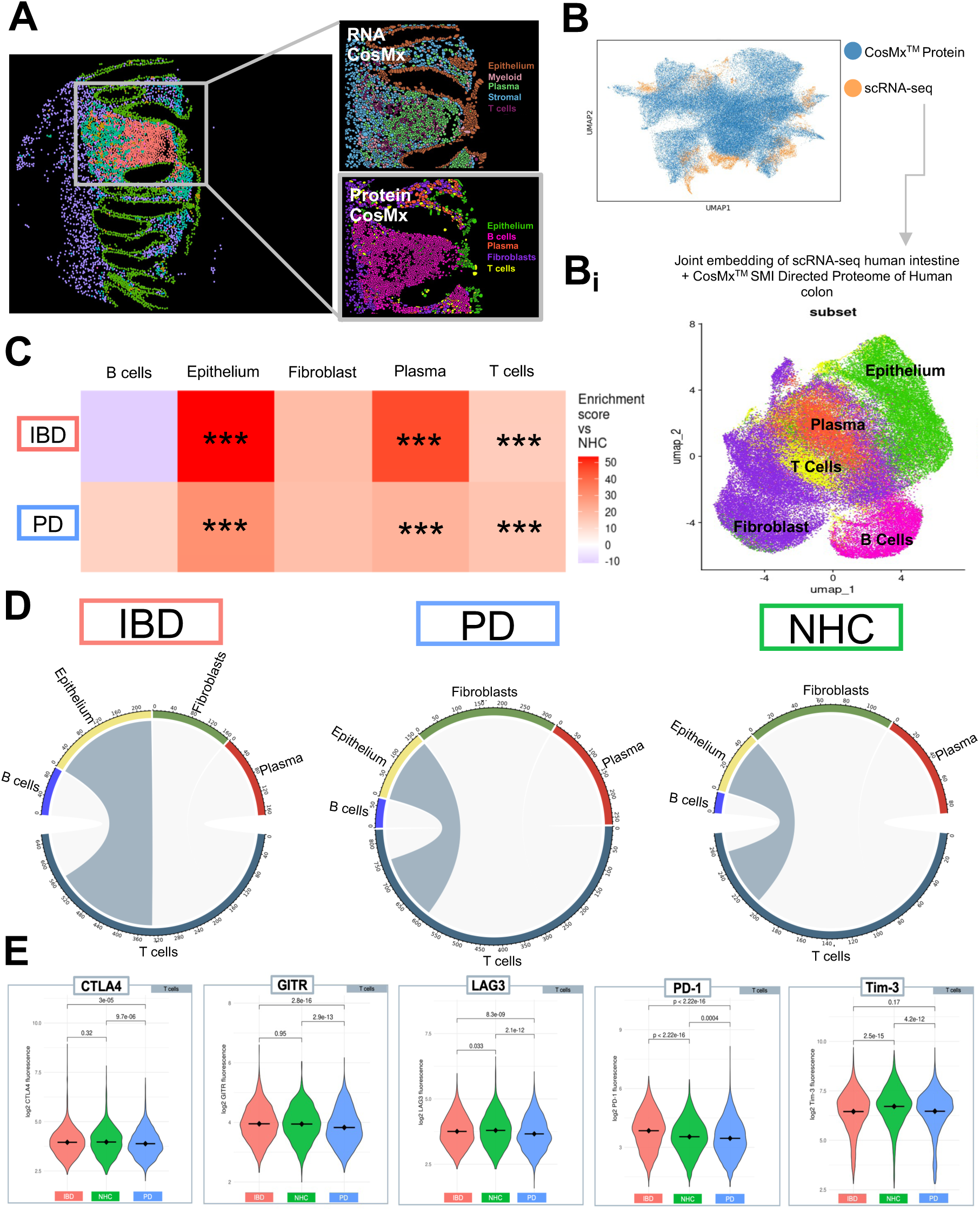
Integration of CosMx^TM^ SMI proteomic and transcriptomic profiles reveals alterations in epithelial cell-T cell communication. **A** CosMx^TM^ protein and CosMx^TM^ RNA images depicting spatial distribution of all annotated subpopulations **B** Uniform manifold approximation and projection (UMAP) of individual cell distribution and dataset overlap between scRNA-seq (Dataset obtained from Garrido-Trigo et al., 2023) and protein dataset**. Bi** UMAP combining scRNA-seq and CosMx SMI data to link gene expression to protein regulation and annotate cell types. Gene and protein datasets were linked using MaxFuse. **C** Heatmap depicting cell-type abundance by IBD or PD as compared to NHC. Statistical analysis completed by Chi-Square (χ²) test with p value adjusted using Benjamini–Yekutieli. Only results with a fold change greater than 1.5 or less than 0.66 are considered significant after adjustment. * = adjusted p-value less than 0.05, ** = adjusted p-value less than 0.01, ***= adjusted p-value less than 0.001. **D** T cells (gray) are the source, and epithelial cells are the target based on inflammatory disease presentation. Chord plots generated via SCOTIA analysis. **E** Violin plots comparing raw fluorescence of individual proteins expressed in T cells, parsed by disease group. Violin plot significance values are generated via Wilcoxon test with Bonferroni p-value correction.

## Discussion

It is known that the gut microbiome and health of the gastrointestinal tract directly shapes the circulating immune system^16^. Peripheral immune dysfunction influences BBB permeability and can thereby compromise brain function via alterations that include chemotactic signaling^20,31,32^. Individuals living with PD commonly and consistently report GI dysfunction decades prior to onset of motor symptoms^2^, as well as have been reported to have a dysbiotic gut microbiome distinct to PD^32–35^. IBD is a commonly occurring condition in the US^13^ which catalyzes chronic systemic inflammation and has been epidemiologically linked to PD^36,37,38^. To investigate the molecular underpinnings of this association, we analyzed the microenvironments of the colonic luminal lining, peripheral blood and stool from individuals with IBD in remission, or with idiopathic PD, or in neurologically healthy controls (NHC) to determine the extent of shared versus unique pathways that could shed light on potential molecular mechanisms linking mid-life chronic gut dysfunction arising from IBD to the increased risk for development of PD later in life.

First, single-cell targeted transcriptomic profiles of sigmoid colonic biopsies revealed cell type-specific phenotypes unique to PD and IBD. Second, differential gene expression analyses of these clusters identified inflammatory pathways associated with immune signaling dysfunction in both PD and IBD colonic biopsies, as compared to NHC. Third, spatial transcriptomics analysis of these colonic biopsies revealed dysregulation in cell-cell crosstalk and neighborhoods in both IBD and PD, that importantly were confirmed and validated at the protein level by proteomic analysis. Finally, multiplexed inflammatory factor immunoassays revealed a dysfunctional immune phenotype in both IBD (stool) and PD (plasma) cohorts, where CCL22 correlated with both PD disease duration and ferritin load, as compared to NHC. Together with literature describing alterations in the abundance of circulating adaptive immune cells in those living with IBD^39^ or PD^40,41^ and shared depletion of key SCFA-producing gut bacteria in IBD and PD^42^, our novel findings suggest that a subset of individuals living with IBD may be at increased risk for PD pathogenesis due to chronic systemic inflammation, likely initiated in the GI tract due to gut dysbiosis and loss of SCFA-producing microbes. Specifically, the chronic peripheral (blood and gut) inflammation in IBD would be expected to impair the ability of the peripheral immune system to mount proper responses to antigenic challenges over time which is a critical requirement to distinguish between self and non-self and maintain immune tolerance. It is well-known that inflammatory challenges *in vivo* hasten upregulation of α-synuclein in various cell types^43–46^; and in the case of α-synuclein-expressing enteroendocrine cells, increasing the likelihood of transfer and propagation to enteric ganglia and vagal efferent connections where it becomes pathogenic^47^.

Although the mucosal lining is the largest immune-antigen interface in the body^48^, to our knowledge this is the first single-cell spatial resolution investigation of the colonic immune mosaic from those living with PD or IBD to look for mechanism-based clues to support the gut-first hypothesis of PD and explain the epidemiological association between these conditions. As a result of GI injury and resulting intestinal epithelial breakdown, up to 79% of IBD patients 18-25 years of age and 90-100% of patients >65 years of age experience iron dysregulation.^49^ Iron dysregulation is also observed in PD, where total iron content in the brain often increases^50–52^ and a decrease in ferritin is observed in the blood^53,54^ of those living with PD. These data suggested to us that iron mishandling may also link these two chronic systemic diseases of inflammation^55^. To our knowledge, there has yet to be any investigation of iron handling in the gut of individuals living with PD. Therefore, we herein further characterized this mechanism at the only anatomical site of iron absorption—the gut mucosal lining. Additionally, a recently published study identified enrichment of ferroptotic pathways in gut-innervating neurons upon inflammatory stimuli that mimic cholitis^56^. Our findings revealed disease-specific differences in transcript accumulation for *FTH1*, the ferritin heavy chain gene critical for primary iron storage, within several immune cell types (Fig. 2E); specifically, we detected significantly increased transcript accumulation of *FTH1* in epithelial cells of IBD patients and significantly decreased in epithelial cells of PD patients compared to NHC (Fig. 2A & B). We also observed the largest difference in *FTH1* accumulation in colonocytes, a subtype of epithelial cells (Fig. 2). We speculate this finding may be related to the inflammatory peripheral stress response associated with PD, which can directly support systemic iron withdrawal by limiting dietary iron absorption^57^. Additionally, *SlLC40A1* which encodes for ferroportin, the primary iron exporter which releases iron into circulation for systemic use, was significantly down regulated in colonocytes from both those living with IBD and PD, as compared to NHC (Fig. 2F). At the single-cell level, we found that those living with IBD (in remission) and PD had significantly more *FTH1+* colonocytes interacting with other cell types, in particular IgA-producing plasma cells, relative to NHC colonocytes which primarily interacted with myofibroblasts (Fig. 4A; Fig. 4C-Ci). Additionally, colonocytes that were *FTH1*+ in those living with IBD in remission displayed significant depletion in *MTOR* and *LGALS* transcripts (Fig. 4D), while colonocytes that were *FTH1+* in those living with PD displayed a substantial enrichment in *MHC I*, as compared to *FTH1*+ colonocytes from NHC. MHC I expression is critical for adaptive immune cell recruitment, specifically cytotoxic CD8 T-cell differentiation^49^. It is known that constant antigen exposure via overexpression of the MHC I complex can drive dysregulation in T cells^58–60^. Together, these findings support a model in which iron mishandling at level of the mucosal lining in colonocytes is a likely catalyst of peripheral (blood and gut) immune dysfunction evident in both IBD and PD, as measured by downregulation of key regulatory immune response genes in a colonocyte subpopulation that is actively transcribing ferritin genes to deal with iron overload, dynamically increasing interactions with IgA-producing plasma cells, and upregulating antigen-presenting complexes that induce cytotoxic T cell differentiation. In parallel with the dysregulated immune phenotype displayed in subsets of colonocytes by spatial transcriptomics, gene ontology (GO) pathway analyses revealed both shared and distinct features in IBD versus PD relative to NHC. Specifically, while there was shared upregulation of lipid biosynthetic processes in IBD and PD colonocytes, IBD colonocytes displayed upregulation of pathways involved in negative regulation of immune and inflammatory responses relative to NHC, while PD colonocytes displayed downregulation of the same processes relative to NHC (Fig.3).

Chronic inflammation associated with tumor microenvironments triggers immune cell dysfunction—where immune cell function greatly decreases, and over time cells become unresponsive to invading pathogens and other stimuli^61^. Therefore, we posit that chronic gut inflammation associated with repeated infections or exposures to environmental risk factors that promote a leaky and inflamed gut may similarly trigger chronic peripheral inflammation and hasten the onset or increase the risk for PD for a subset of individuals with IBD. We identified a depletion in a key T-cell recruitment chemokine CCL22 in IBD stool and PD plasma, as compared to NHC (Fig. 5). The decrease in CCL22 expression may be the result of dysregulation of immune-related proteins in colonic T cell populations in IBD (in remission) and PD, as compared to NHC (Fig. 6E). These markers of reduced immune competence may contribute to increased susceptibility to the effects of chronic infections and risk for age-related disease^62,63,64,65^. Adaptive immune cells rely on iron metabolism to perform effector functions— such as proliferation and migration^66^. Circulating T cells from the blood of individuals living with PD have been reported to display decreased migratory capacity and proliferative phenotypes^67^. Interestingly, iron-deficient mice display a reduction in T cell proliferation and an impaired immune response that is not recapitulated in mice experiencing iron overload^68^. Furthermore, iron dyshomeostasis can promote a leaky gut as well as impair immune function, this may account for the phenotype of immune signaling dysfunction within T cells we observed near the epithelial barrier (Fig. 6C-E). It is known that chronic systemic inflammation increases the risk of PD^64^. One possible hypothesis is the downregulation of the adaptive immune response, which could be a protective peripheral mechanism to deplete a chronic cytotoxic inflammatory response.

In summary, it is known that the clinical manifestations of PD can remain dormant for decades and appear long after non-motor symptoms start. We speculate that chronic immune activation in the gut due to gut dysbiosis, invading pathogens, pesticide/toxicant exposure, IBD, and/or diets rich in carbohydrates represent environmental triggers that synergize in a subset of individuals to promote a leaky gut. This can induce the dampening of immune response and is further catalyzed by iron mishandling, which can result in inappropriate immune signaling along the gut-brain axis in individuals at risk for development of PD^69^. Together, our data support a model in which sustained chronic inflammation in several peripheral compartments likely converges and creates the perfect storm that accelerates the transition from a prodromal pre-clinical premotor stage of PD to a clinical motor stage. Additional research will be needed to reproduce and validate our findings and the potential utility of gut immune phenotyping to identify individuals at high risk for development and/or progression of PD who can be enrolled in secondary prevention trials. These foundational data set the stage for future investigation and therapeutic discovery that targets the integrity of the gut mucosal lining to modulate chronic systemic inflammatory cascades that are likely to increase the risk for PD development and progression.

## Methods

### Participant recruitment

The University of Florida Institutional Review Board (IRB201902459) reviewed and approved this study. Individuals living with PD were recruited from the University of Florida Neuromedicine Clinic at the Fixel Institute for Neurological Diseases. Individuals living with IBD who were already preparing for routine colonoscopy were recruited from the UF Gastrointestinal (GI) Clinic. All participants consented to this study and signed an informed consent document prior to sample collection. Neurologically healthy control I(NHC) individuals were all PD-spousal controls except for one subject that was an IBD spousal control.

Participation within this cross-sectional study included inclusion criteria of reported age between 40-80 years, diagnosis of PD (based on MDS score), IBD (ulcerative colitis and/or Crohn’s disease specified if applicable), or no diagnosis in the case of the neurologically healthy control (NHC) cohort. Individuals were excluded if individuals living with PD reported use of immunosuppressants for active infections or active autoimmune or chronic inflammatory conditions. Individuals were also excluded if they reported a history of antibiotic use within a month or recruitment. Additional exclusion criteria applicable included if an individual reported being pregnant, if the individual reported a history of a blood transfusion within 4-weeks of recruitment, reported body weight less than 110 lbs. Additional health history collected included: assessment of disease duration, MoCA and Schwab scores, past or current endocrine dysfunction, and cancer history, age, sex, race, lifestyle habits such as smoking, over-the-counter or prescription NSAID use. Primary selection criteria for this cohort included age then sex matching as close to the NHC as possible, therefore due to cost and assay size not all patients who provided biopsies were used within this series of assays. Despite intense efforts toward recruiting a diverse population of participants, we were unable to obtain quality representation of non-white ethnic groups within our study. Future studies will focus on recruitment of a more diverse population. These metadata were collected from all participants and anonymized data are available upon request. Of importance and unique to this cohort, all IBD and PD participants within this study were in endoscopic remission (not actively inflamed at the level of the gut), confirmed by UF GI clinic gastroenterologists.

### Sample collection and preparation

Sigmoid colonic biopsies were collected via sigmoidoscopy in individuals with PD or NHC and via colonoscopy in individuals with IBD by a board-certified UF gastroenterologist via protocols approved by the UF IRB and fixed in formalin and paraffin embedded (FFPE) to be used for CosMx^TM^ SMI. Within these data, *N* = 12 PD biopsies, *N* = 13 IBD biopsies, *N* = 9 NHC biopsies. Two 5mm FOVs were drawn per biopsy, based on H&E of subsequent slide as well as aiming to obtain the most surface area of the total core. In brief, FFPE sample preparation for imaging included punching 1mm tissue microarray cores from each patient sample block to generate a tissue microarray (TMA) at 5µm section thickness and allow for all samples to be ran on one slide in the same run with the same lot numbered reagents. A more detailed description of the protocol designed by the Tansey lab for TMA production using FFPE prepared tissue may be found on protocols.io at URL: dx.doi.org/10.17504/protocols.io.5jyl8drqrg2w/v1. Blood and stool samples were also collected from this participant cohort for protein analysis. All protocols, software/code, and datasets may be obtained via the key resource table attached at the final supplementary Table 3 via excel.

### Spatial Molecular Imaging (SMI) analysis: targeted transcript and proteomics CosMx^TM^ SMI: directed transcriptomics and proteomics

All protocols for single-cell spatial workflow were followed exactly to manufacturer instruction with no adaptation, as described in CosMx*^TM^* manual provided, as well as^12^. Via the use of multiplexed capture probes and reporter probes, NanoString CosMx*^TM^*can spatially resolve up to 1000 transcripts (CMx Hs Univ Cell Panel RNA Kit EA: CAT# 121500002) and 64 proteins (CMx Hs IO Panel Protein Kit: CAT# 121500010). A custom spike in for the following 12 genes labeled following the HUGO gene nomenclature were included in the RNA assay: *SLC6A4* (solute carrier family 6 member 4), *CHGA (chromogranin A)*, *SLC11A2 (solute carrier family 11 member 2)*, *FTH1(ferritin heavy chain 1)*, *HAMP (hepcidin antimicrobial peptide)*, *IREB2 (iron responsive element binding protein 2)*, *ITPKB* (Inositol-Trisphosphate 3-Kinase B), *NDUFB1* (NADH:Ubiquinone Oxidoreductase Subunit B1), *PINK1* (PTEN-induced kinase 1), *PYY* (peptide YY), *RAB8A* (RAB8A, member RAS oncogene family), *TF* (transferrin). It is important to note that due to assay restrictions, the protein sections are 5 µM from RNA sections and thus were run on separate slides. We used morphology markers CD45, PanCK, CD3 to visualize the tissue and picked two 0.5mm fields of view (FOV) per sample. To obtain cell count matrices for each slide, cell boundaries were generated via morphology antibodies described above and localization of transcript expression, as previously described^70^. Individual analysis pipelines for either RNA or protein are described below.

### Quality control and processing of CosMx^TM^ RNA

Cells were flagged if: the cell had less than 20 counts, if 10% of the counts were negative probes, total counts did not exceed the number of detected genes, the cell had less than 10 features, the cells had area outliers based on Grubb’s test p-value(0.01) and the cell had a mean of 0.5 negative counts. Flagged cells were then removed from downstream analysis. An average of 89.29% cells passed quality control (QC). After QC, a total of 75,170 cells were used in downstream analysis. Then SC Transform (Seurat package, version 5.1.0) was leveraged for normalization.

### Cell-type assessment and annotation for CosMx^TM^ RNA

Supervised clustering was preformed using an existing scRNA-seq intestine database^12^ as a reference. Immunoglobulins (Igs) were removed from downstream analysis due to high expression within all cell-types. Cell subset classification was then preformed using immunofluorescence data obtained from the morphology markers described above (PanCK, CD45, Mean. DAPI, mean CD68) as well as gene expression, and the reference scRNA-seq object using InSituType (InSituType version 1.0.0). Supervised clustering then generated 5 primary clusters: epithelial, myeloid, plasma, stroma and T cell. Cells with 75% or more probability to be that specific subset were used for downstream analysis. Within each subset, the same clustering methodology was performed to identify refined (i.e., from T cells parsing CD4 and CD8).

### Quality control and processing of CosMx^TM^ Protein

Cells were used in downstream analysis if 50% or more of the proteins were in the 90^th^ percentile or higher of expression. Additionally, cells were removed from downstream analysis if the cell contained less than 10 proteins that were also in the 50^th^ percentile or lower of expression. Cells were also removed from downstream analysis if the cells negative probe threshold was lower than 2 and above the upper threshold of 15 and/or contained area outliers based on Grubb’s test p-value (0.01). After QC, 51.25% of cells were kept, which resulted in 77,057 total cells. For cell normalization, average total intensity scaling and arsinh transformation with a cofactor of 50 were implemented.

### Cell type assessment and annotation for CosMx^TM^ Protein

Due to the size of the CosMx™ Human Protein Immuno-Oncology (63 total proteins) panel and to provide information from multiple biological modalities, we used MaxFuse^30^ to integrate an existing scRNA-seq dataset of the intestinal mucosa from NHC^12^ to most accurately annotate cell types. Annotations were then curated by using only the weak links between the 63 proteins and genes present in the scRNA-seq dataset (Supplemental Fig. 4). These data as well as detailed annotation methods are available at https://servidor2-cibrehd.upc.es/external/garrido/methods1/.

### Abundance analysis

Aggregation of total cell counts per patient were leveraged to identify abundance of cell type by cluster by health status (Fig. 1E, RNA; Fig. 6C, protein). Cluster frequencies were compared using Chi-squared tests (X^2^) for enrichment and Wilcoxon rank-sum test to account for patient variability. Fold changes were calculated, and p-values were adjusted via the Benajmmini Yekutieli method (adj p <0.05) where significant clusters were annotated with a FC > 1.5 or < 0.66 (Fig. 6C).

### Cell-to-cell interaction

CellTalkDB, accessed via https://github.com/ZJUFanLab/CellTalkDB, to understand ligand receptor interactions. Then the SCOTIA python package, accessed via https://github.com/Caochris/SCOTIA?tab=readme-ov-file, was used to obtain cell-cell communication by disease (Fig. 4B,D,F RNA; Fig. 6D protein).

### Multiplex immunoassays on Meso Scale Discovery (MSD) Platform

Protein was isolated from stool samples using MSD lysis buffer (MSD; R60TX-3), 1 tablet complete protease inhibitor (Roche) and 5 mm stainless steel bead (Qiagen). After solids were removed by separating protein supernatant, protein concentration was determined via BCA Protein Assay Kit (Pierce), according to manufacturer instruction. Duplicates of stool supernatant (25ul) were diluted 1:1 and used to quantify chemokine and cytokines via U-PLEX custom pro-inflammatory human panel (Eotaxin, Eotaxin-3, IFN-γ, IL-1β, IL-6, IL-8, CCL22 (MDC), TNF-α (MSD; K15067M-1) and R-PLEX human ferritin (MSD; F21ADA-3) on the Quickplex MSD instrument, according to the manufacturer’s protocol.

### Data availability

All processed CosMx*^TM^* SMI flow cell data may be explored via Zenodo online data repository URL: via Zenodo: https://doi.org/10.5281/zenodo.14851478 (RNA) https://doi.org/10.5281/zenodo.14851272 (protein).

### Code availability

All code used for the analysis and plot is available at https://github.com/ibd-bcn/CosMx-Bolen. All Code for scRNAseq used for cell phenotyping is available at <u>1</u> or on Salas lab website at https://servidor2-ciberehd.upc.es/external/garrido/methods2/.

## Acknowledgements

The Tansey lab and all collaborators would like to acknowledge and thank all participants within this study. Additionally, thank you to Dr. Rebecca L. Wallings and Dr. Kelly B. Menees who provided insightful feedback and guidance on this manuscript. We would also like to highlight that this work is the wonderful outcome of global team science. Due to Dr. Tansey’s focus on collaborative science that pushes the frontier of medicine, we have built a dynamic team and generated datasets that would have been impossible without multidisciplinary crosstalk across several institutions.

---

**Supplemental Figure 1.**
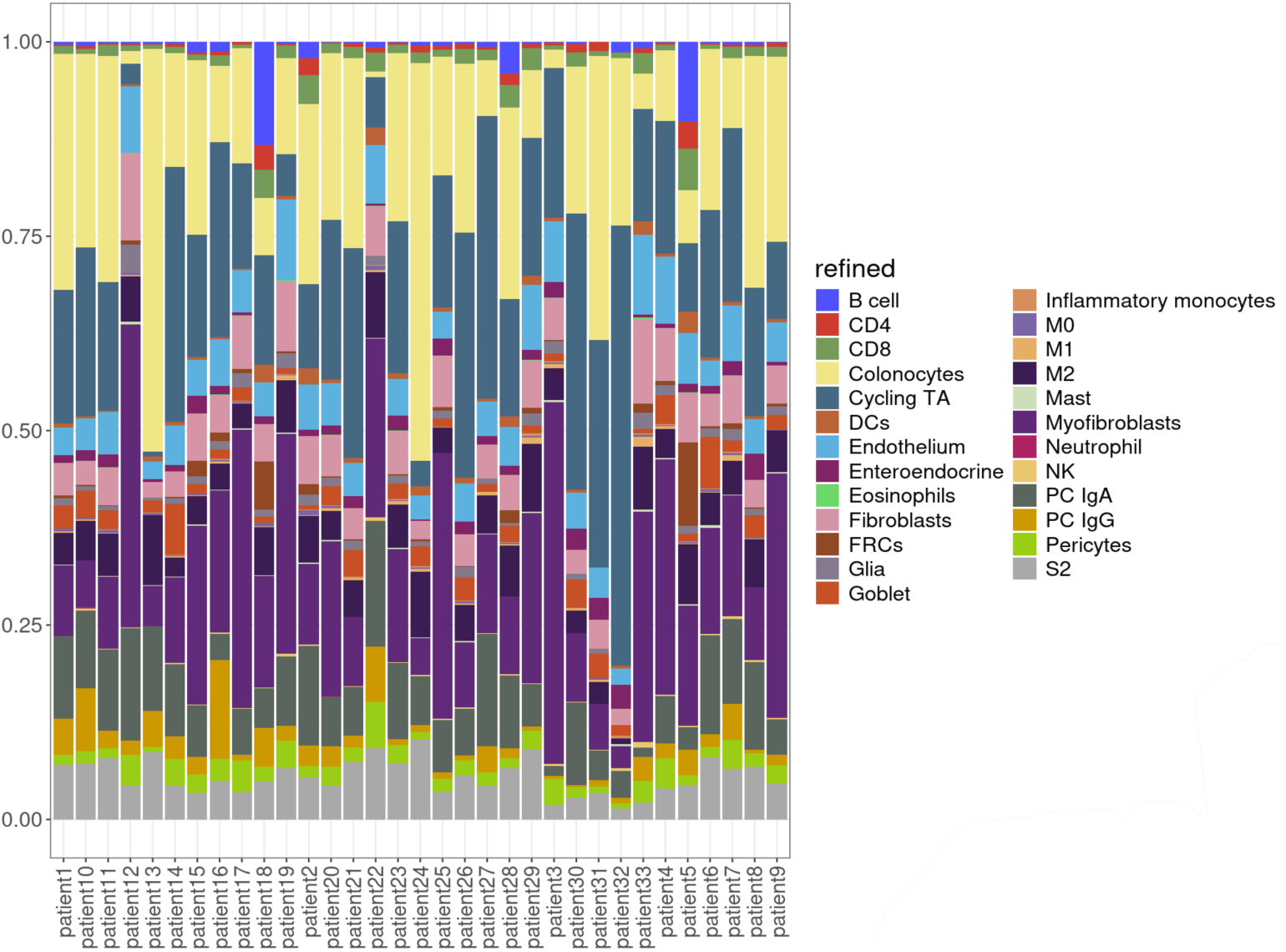
Refined cell-type abundance from every participant.

**Supplemental Figure 3.**
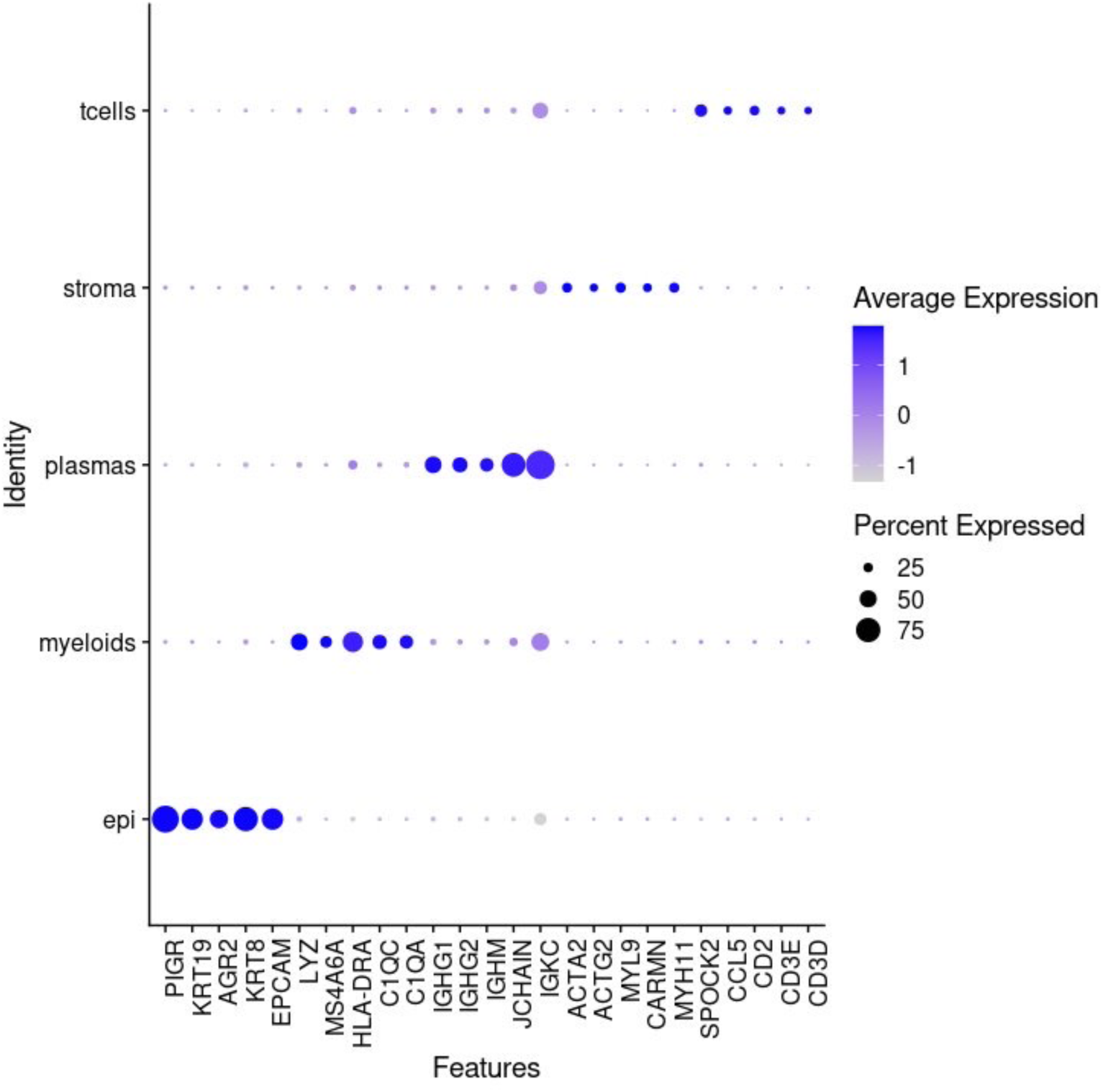
CosMX SMI transcriptomics Cell cluster identifiers.

**Supplemental Figure 4.**
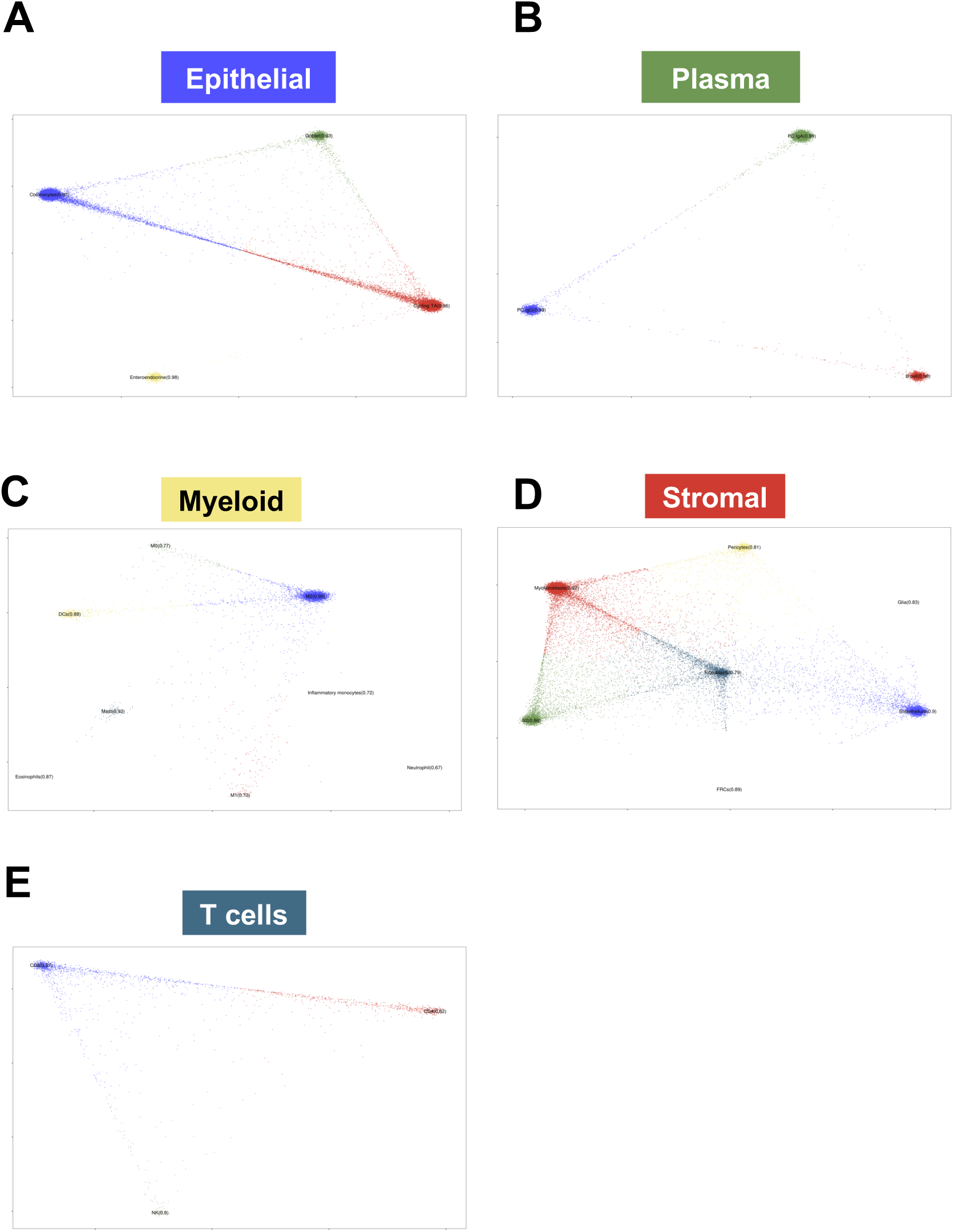
CosMx SMI refined gene cluster flight-path analysis.

**Supplemental Figure 5.**
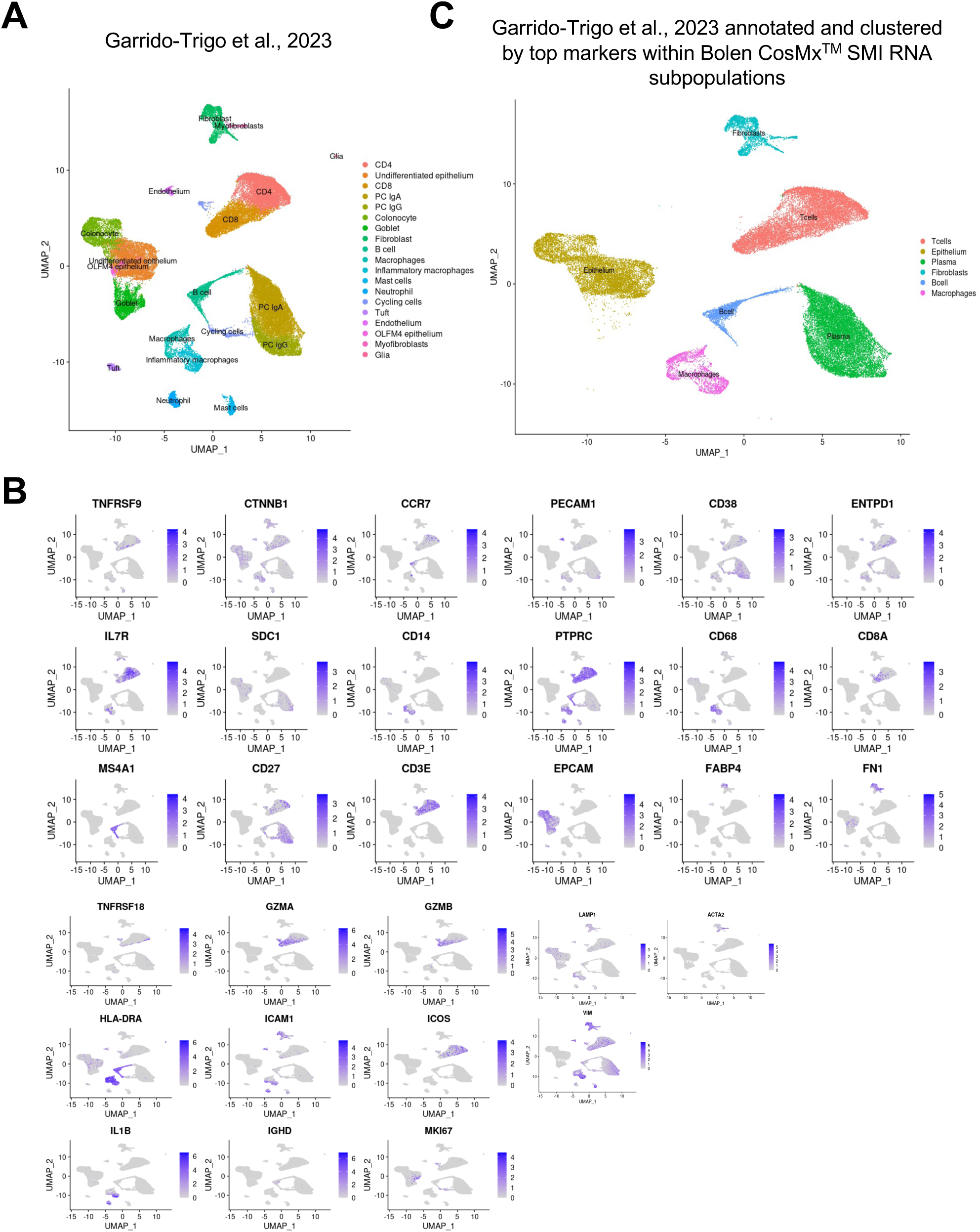
Single-cell targeted proteomic analysis successfully integrates with scRNA-seq and depicts multimodal immune dysregulation across multiple biological outputs of human colonic biopsies.

**Supplemental Figure 6.**
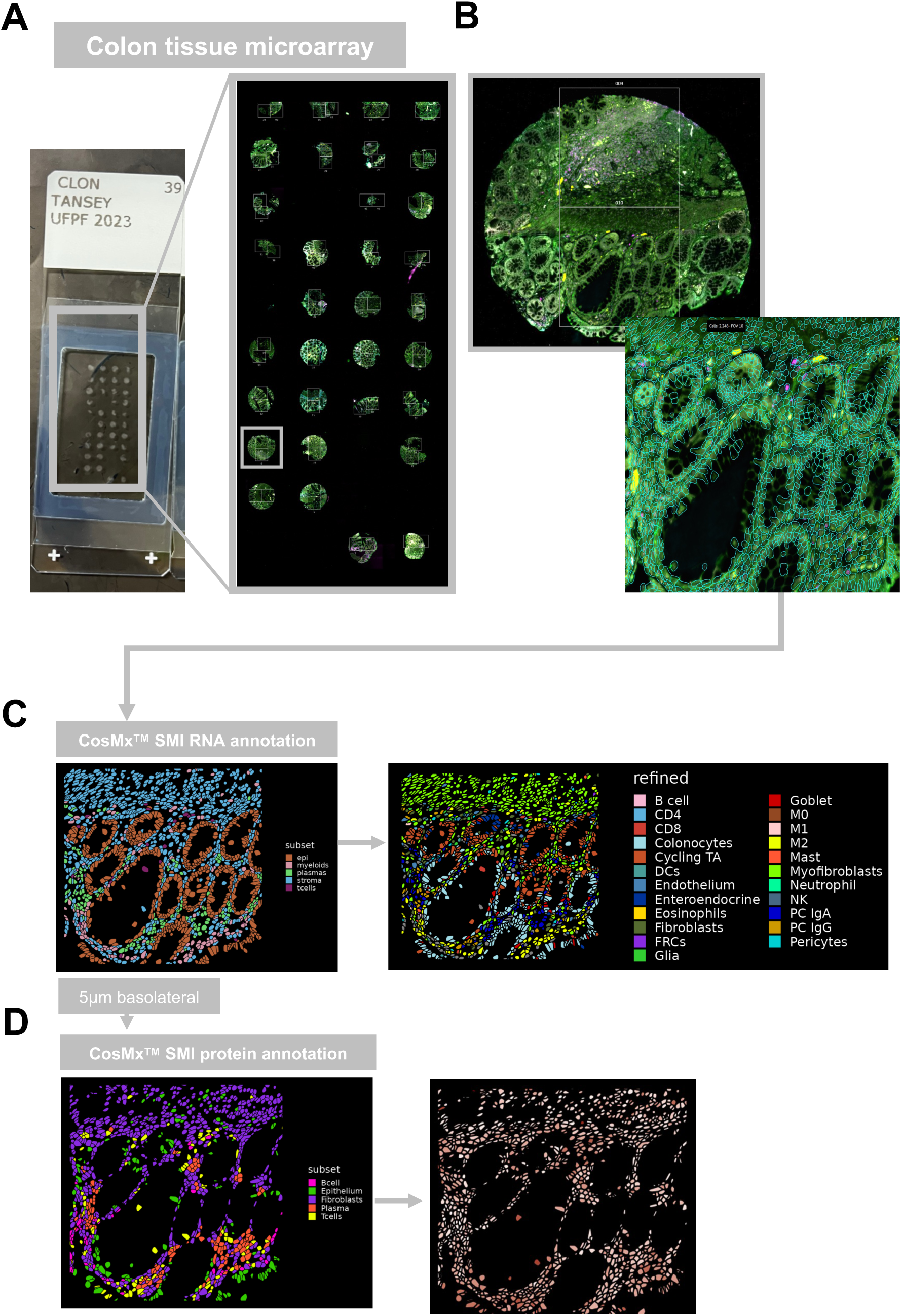
Analysis workflow of CosMx^TM^ SMI RNA and protein analyses.

**Supplemental Figure 7.**
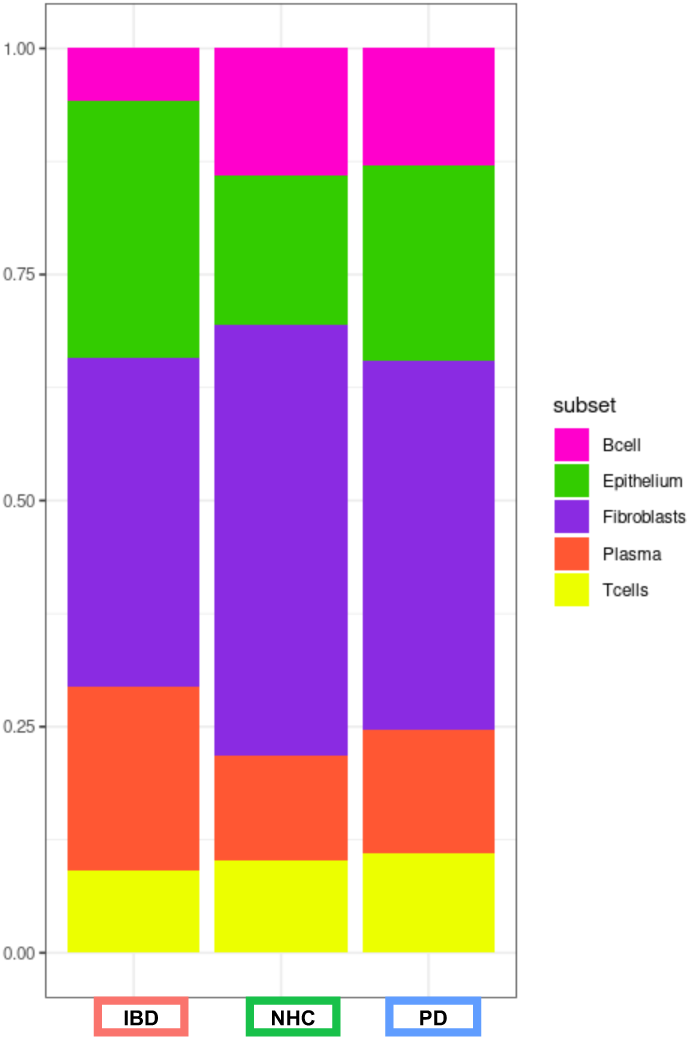
Spatial molecular imaging (SMI) of directed proteome provides cell-type specificity and frequency at the level of subpopulations in sigmoid colon biopsies from those living with inflammatory bowel disease in remission (IBD) and Parkinson’s disease (PD).

**Supplemental Table 1.**
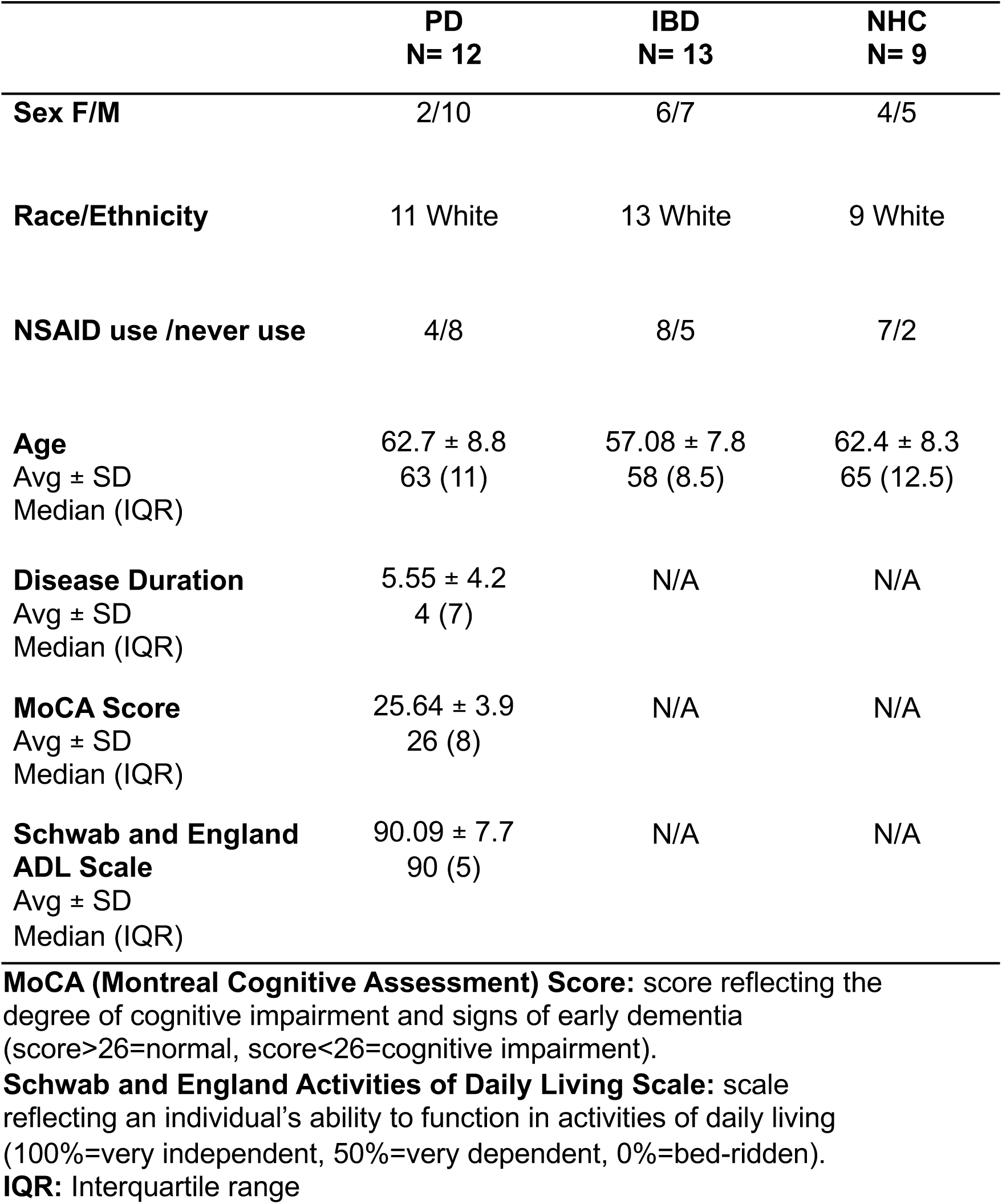
Colonic biopsy patient demographics and cognitive status.

**Supplemental Table 2.** CosMx SMI gene cluster identifiers.

| Cell-type | Subpopulation Markers | Refined population markers |
| --- | --- | --- |
| Epithelial cells | EPCAM, AQP8, BEST4, MUC2, OLFM4, PLCG2, TRPM5, ZG16 | Colonocytes: AQP8, FABP1 and SLC26A2<br>Goblet cells: MUC2, TFF3 and SPINK4<br>Enteroendocrine: CHGA<br>Cycling TA: MKI67, TOP2A and PCNA |
| B cells | CD79A, BANK1, CD19, DERL3, MS4A1, MZB1 | NA |
| Plasma cells | DERL3, MZB1, XBP1 | PC IgA: +IgA and DERL3, MZB1, XBP1, -IgG<br>PC IgG: +IgG and DERL3, MZB1, XBP1, -IgA |
| T cells | CD3D, CD3E, CD3G, CD8A, FOXP3, GZMA, GZMB, IL17A, NKG7, TRBC1 | CD8: CD8A, CD8B, GZMK, KLRB1, SPRY1 and IGTA1 and KLRG1<br>CD4: CD4, CCR7, LEF1, ANXA1, IL7R and GPR186<br>and SELL<br>NK: KLRF1, NCAM1, and lacks expression of CD3G, CD3D, CD8A and CD4 |
| Stromal cells | ACTA2, ADAMDEC1, CHI3L1, COL3A1, NRXN1, PVALP, SOX6, VWF | S2: VSTM2, NPY and NRG1<br>Endothelium: VWF, PECAM1 and PLVAP<br>Fibroblasts: ADAMDEC1, CP, OGN, CCDC80 and GREM1<br>and FABP4<br>FRCs: CCL19 and CCL21<br>Glia: NRXN1, S100B and CDH19<br>Myofibroblast: SOSTDC1, ACTG2 and MYH11<br>Pericyte: NOTCH3 and RGS5 |
| Myeloid cells | AIF1, C1QA, C1QB, CD14, CMTM2, FCGR3B, LYZ, MS4A2, TPSAB1, TPSAB2 | M0: lack the expression of M2 markers and expression of CD68, C1QA, C1QB, SELENOP, LYZ, HLA-DPB1, HLA-DPA1, AIF1<br>M1: ACOD1, TNIP3, IL1B, INHBA, IL6, VCAN, CD300E, CXCL5, TNIP3, IL1B, INHBA, IL6, VCAN, CD300E<br>M2: CD209, CD163L1 and FOLR2<br>Inflammatory monocytes: VCAN, CD300E, CD14, FCN1 and S100A9<br>Mast cells: LTC4S, TPSB2 and TPSAB1<br>Eosinophils: CLC, IL4 and IL13<br>Neutrophils: PROK2, CMTM2, CXCL8, FCGR3B, AQP9, S100A8 and S100A9<br>DCs: CD1C, TRL10, TCTN3<br>CCL22, CCL19, LAMP3 |

